# An efficient method for viable cryopreservation of hookworms and other gastrointestinal nematodes in the laboratory

**DOI:** 10.1101/2023.02.01.526637

**Authors:** Hanchen Li, David Gazzola, Yan Hu, Raffi V. Aroian

## Abstract

Hookworms (genera *Ancylostoma* and *Necator*) are amongst of the most prevalent and important parasites of humans globally. These intestinal parasites ingest blood, resulting in anemia, growth stunting, malnutrition, and adverse pregnancy outcomes. They are also critical parasites of dogs and other animals. In addition, hookworms and hookworm products are being explored for their use in treatment of autoimmune and inflammatory diseases. There is thus a significant and growing interest in these mammalian host-obligate parasites. Laboratory research is hampered by the lack of good means of cryopreservation. Here, we describe a robust method for long-term (≥3 year) cryoprotection and recovery of both *Ancylostoma* and *Necator* hookworms that is also applicable to two other intestinal parasites that passages through the infective third larval stage, *Strongyloides ratti* and H*eligmosomoides polygyrus bakeri*. The key is the use cryo-preserved first-staged larvae raised to the infective third larval stage using activated charcoal mixed with uninfected feces from a permissive host. This technique will greatly facilitate research on and availability of gastrointestinal parasitic nematodes with great importance to global health, companion animal health, and autoimmune/inflammatory disease therapies.

## 1. INTRODUCTION

Hookworms (*Necator americanus, Ancylostoma ceylanicum*, and *Ancylostoma duodenale*) are blood-feeding helminth parasites of humans that live in the small intestine and are responsible for >4 million disability-adjusted life years (DALY) and $100 billion of economic losses annually (Jourdan et al. 2018; Loukas et al. 2016; Pullan et al. 2014; Umbrello et al. 2021; Bartsch et al. 2016; Hamory et al. 2021). Hookworm infection has had a significant influence on human history and is a major cause of morbidity in infants, children, and women of child-bearing age in the tropics and subtropics leading to, amongst other things, malnutrition, permanent growth stunting, cognitive impairment, adverse pregnancy outcomes, decreased worker productivity, and decreased educational status and future earnings. The closely related hookworm *Ancylostoma caninum* is the most prevalent and important intestinal nematode parasite of dogs in the United States of America (Jimenez Castro et al. 2021, 2020). Its prevalence is rapidly spreading, along with a multidrug resistant phenotype, making it increasingly recalcitrant to treatment (Jimenez Castro et al. 2021, 2020). Development of new and better therapies and a protective hookworm vaccine are considered important goals for global and animal health (Loukas, Maizels, and Hotez 2021; Haldeman, Nolan, and Ng’habi 2020; Shepherd, Wangchuk, and Loukas 2018; Mourão Dias Magalhães et al. 2021; Hu, Nguyen, et al. 2018; Noon et al. 2019).

Hookworms and other helminth parasites are also major drivers in the evolution of the vertebrate and human immune systems, in particular in shaping the Th2 immune response (Fumagalli et al. 2010; Maizels et al. 2009; Loukas et al. 2016). The pressure imposed by parasitic worms on human genes has been hypothesized to be stronger than that of viral, protozoa, or bacterial agents (Fumagalli et al. 2011). Because of their strong immunomodulatory characteristic, hookworms, most notably *Necator americanus*, and their secreted products are garnering great interest for treatment of autoimmune and autoinflammatory diseases such as celiac disease, asthma, metabolic syndrome, multiple sclerosis, and inflammatory bowel diseases (Loukas, Maizels, and Hotez 2021; Mourão Dias Magalhães et al. 2021; Loukas et al. 2016; Montaño, Cuéllar, and Sotillo 2021; Ryan et al. 2020; Chapman et al. 2021).

Studies on human hookworms are therefore of enormous and growing interest vis-a-vis global health, drug development, vaccine development, immunology, and autoimmune research and therapies. Currently, *Necator* and/or *Ancylostoma* hookworms can be maintained and studied in the laboratory using hamster, dogs, and even humans as hosts (Loukas et al. 2016; Chapman et al. 2021; Montaño, Cuéllar, and Sotillo 2021). There are, however, significant challenges in working with these obligate mammalian parasites in the laboratory that are exacerbated by the lack of reliable cryopreservation methods (see Discussion). For example, laboratories working with these parasites are under constant pressure to maintain their hookworm lines or else permanently lose their invaluable cultures. Methods for long-term preservation of hookworms has thus been deemed critically important for continued human experimental infection and therapy (Chapman et al. 2021). Here, we describe methods for the long-term (≥3 year) cryoprotection and recovery of viable *Necator* and *Ancylostoma* hookworms as well as other gastrointestinal parasitic nematodes that infect via infectious third stage larvae (iL3).

## 2. MATERIALS AND METHODS

### Medium and reagents

Cryprotectant solution was made similar to as described (Duarte, Harrison, and Cappello 2003): 70% RPMI 1640 (Gibco Cat# 11835-030), 10% dimethyl sulfoxide, 10% dextran T10 (Pharmacosmos A/S, DK-4300 Holbaek, Denmark), and 10% Fetal Bovine Serum. Dexamethasone (DEX) was used to immune suppress the hamsters for *N. americanus*-related experiments (Hu, Nguyen, et al. 2018). S Medium was prepared as described (Sulston and Hodgkin 1988). Images were taken with an Olympus SZ-CTV dissecting microscope fitted with an Infinity1 camera. GraphPad Prism v. 9 was used for all graphs and analyses.

### Animals and parasites

*A. ceylanicum* and *N. americanus* hookworms were maintained as previously reported (Hu, Nguyen, et al. 2018; Hu et al. 2012; Li et al. 2021). Three to four-week-old male Golden Syrian hamsters (HsdHan:AURA) were purchased from Envigo (U.S.A) and were infected at approximately 4–5 weeks of age with either ∼130 *A. ceylanicum* third-stage infectious larvae (iL3) orally or ∼300 *N. americanus* iL3 larvae subcutaneously. Hamsters were provided with food and water (ad libitum). For isolation of iL3s for direct freezing, infected hamster feces was cultured on activated charcoal for 7 days at 28°C and harvested using the Baermann technique.

*S. ratti* was maintained in 6-week old male Wistar rats by subcutaneous injection with 500-1000 iL3s. Infected feces were cultured on activated charcoal for 5-7 days at 22°C to isolate iL3s. *H. polygyrus bakeri* was maintained in 6-week old male Swiss Webster mice by oral gavage with 200 iL3s as previously reported (Hu et al. 2010). Infected feces were cultured on activated charcoal for 7 days at 22°C to isolate iL3s.

All animal experiments were carried out under protocols approved by the University of Massachusetts Chan Medical School (IACUC protocols 202100090, 202000071, 202000044). All housing and care of laboratory animals used in this study conform to the N.I.H. Guide for the Care and Use of Laboratory Animals in Research (see 18-F22) and all requirements and all regulations issued by the USDA, including regulations implementing the Animal Welfare Act (P.L. 89-544) as amended (see 18-F23). Fecal egg counts and small intestinal worm burdens were determined as previously described (Hu et al. 2010; Hu, Nguyen, et al. 2018; Hu et al. 2012). For *S. ratti*, fecal egg counts were taken seven days post-inoculation.

### L1s freezing, recovery, and growth to iL3 stage

The same procedure was carried out for all parasites and all rodent hosts. Feces from nematode-infected rodents were collected and eggs isolated as described (Mes, Eysker, and Ploeger 2007) with slight modifications. Basically, feces were collected and soaked in a beaker with 13% NaCl solution (2 mL per gram of feces) for 30 minutes at room temperature. The feces were then homogenized with a spatula and the resultant solution was poured through a stainless-steel mesh strainer into a 50 mL conical tube to filter out large material. The tube was then spun at 2000g room temperature for 5 minutes (eggs in the supernatant). The supernatant was decanted into a beaker into which an equal volume of distilled water was added. The resultant egg suspension was split into 50 mL conical tubes and spun as above (eggs pellet). After the supernatant was discarded, the pellets were combined and resuspended in 10 mL 17% sucrose solution and transferred to a 15 mL conical tube. The suspension was spun as above (eggs in the supernatant). The supernatant was split into two 15 mL conical tubes, filled to the top with distilled water, and spun as above (eggs will be in the pellet). The pellets were resuspended in 4.5 mL distilled water to which is added 0.5 mL 6% hypochlorite solution (Fisher Scientific Cat# SS290-1) and gently rocked for 1 minute followed by centrifugation (800g for 2 minutes) and three times washes with sterile double distilled water. After the final wash with water, the eggs were resuspended in 25 mL S Medium and filtered with a 70 μm cell strainer to remove the small debris.

The isolated and surface sterilized eggs were transferred to a 50 cm^2^ tissue culture flask and incubated at 28°C for 42 hours to hatch into L1s. The hatched L1s were spun down at 800g for 2 min at room temperature, resuspended in the cryoprotectant solution at the density of about 2 ×10^4^ worms/ml, incubated at room temperature for 1 hr, and then transferred to a 2 mL cryovial (Simport T301-2). The cryovials were then placed into the Nalgene Cryo 1°C Freezing Container (Nalgene Thermo Scientific,Mr. Frosty, Cat# NL-51000001), which was stored at -80 overnight. The next day, all the cryovials moved to liquid nitrogen for long-term storage.

For recovery and growth of frozen L1s, 5-8 g of fresh feces from relevant host (hamsters for hookworms, rats for *S. ratti*, mice for *H. polygyrus bakeri*) were collected in soaked in an equal volume of tap water for 10-20 minutes in a beaker. The feces were mixed using a spatula with an equal amount of activated charcoal (Sigma-Aldrich, Cat#: C2889) and transferred to a 100×25 deep petri dish (USA Scientific Catalog #8609-0625). The frozen L1 larvae were thawed by plunging the cryovial into a 50°C water bath and shaking the tube slowly. The larvae in solution were transferred to a 15 mL conical tube, to which 13 mL of distilled water were added. The larvae were pelleted at 800g for 2 minutes, resuspended in 10 mL tap water, incubated at 28°C for 1 hr, checked for viability, recentrifuged, and then resuspended in 1 mL tap water. The L1s were then loaded onto the top of the fecal-charcoal mixture and incubated at 28°C for 7 days for hookworms and 22°C for 7 days for *S. ratti* and *H. polygyrus bakeri*, at which point the iL3s were collected by the Baermann technique and washed with sterile double distilled water three times. After the last wash, the iL3 worms were resuspend with B.U. Buffer and stored in a 25 cm^2^ tissue culture flask (Genesee Scientific cat#25-207) at room temperature until they were used to infect host rodents as per standard protocols.

#### Infection efficacy studies

*A. ceylanicum*: Male hamsters were infected *per os* with 130 live iL3s for infection efficacy experiments. The hamsters were sacrificed on day 22 post-inoculation (PI) and intestinal parasite burdens and fecal egg counts were determined as described previously (Hu, Nguyen, et al. 2018; Hu et al. 2012, 2013).

*N. americanus*: Male hamsters were infected subcutaneously with 300 live iL3s for infection efficacy experiments. The hamsters were sacrificed on day 55 PI and intestinal parasite burdens and fecal egg counts were determined as described previously (Hu, Nguyen, et al. 2018; Li et al. 2021).

*S. ratti*: Male rats were subcutaneously inoculated with 500 iL3s, and fecal egg counts were taken on day 7 PI.

*H. polygyrus bakeri*: Male mice were inoculated *per os* with 200 iL3s, and fecal egg counts were taken on day 16 PI as described previously (Hu et al. 2010).

## 3. RESULTS

### Cryprotection results with published protocols

An earlier publication noted the freezing of *Ancylostoma duodenale* hookworm first-stage larvae (L1), recovery of live larvae after 6 months, growth to the iL3 stage on sterile agar plates using exogenous *Escherichia coli* (Aikens and Schad 1989), and infection into a single immunosuppressed dog (Nolan et al. 1994). Although L1s from

*N. americanus* hookworms were also frozen in this study, the ability of iL3s derived from these L1s grown on sterile on agar plates with *E. coli* to successfully infect a dog were not tested (Nolan et al. 1994). To test this, we similarly froze *N. americanus* L1s, thawed and grew them in S medium with exogenous *E. coli* for 7 days at 28° C until reaching iL3 (Hu, Miller, et al. 2018), and infecting into two immunosuppressed hamsters with 300 iL3 each (Li et al. 2021; Hu, Nguyen, et al. 2018). Neither infection was successful, despite the apparent health of the iL3. This result suggested to us that this protocol might not be optimum for *Necator* hookworms.

A different approach involving the direct freezing of iL3s was more recently been reported for cryopreservation of exsheathed and non-exsheathed *A. ceylanicum* (Duarte, Harrison, and Cappello 2003). Direct freezing of iL3s is also used for cryopreservation of the sheep gastrointestinal nematode parasite, *Haemonchus contortus* (Chylinski et al. 2015). We used the *A. ceylanicum* cryoprotection protocol (non-exsheathed iL3) to freeze *A. ceylanicum* and *N. americanus* iL3 (Figure 1). When harvested from fresh cultures prior to freezing, virtually all iL3s of *A. ceyalnicum* (Figure 1A) and *N. americanus* (Figure 1B) were curved, active, and alive. However, following freezing, very few (∼1%) of *A. ceylanicum* iL3s appeared alive based on curvature and motility (Figure 1C) and no *N. americanus* iL3s appeared alive (Figure 1D). These results suggested that this freezing protocol was not efficient for *Ancylostoma* and that *Necator* hookworms were even more difficult to successfully cryopreserve (as was also suggested in the study above).

**Figure 1.**
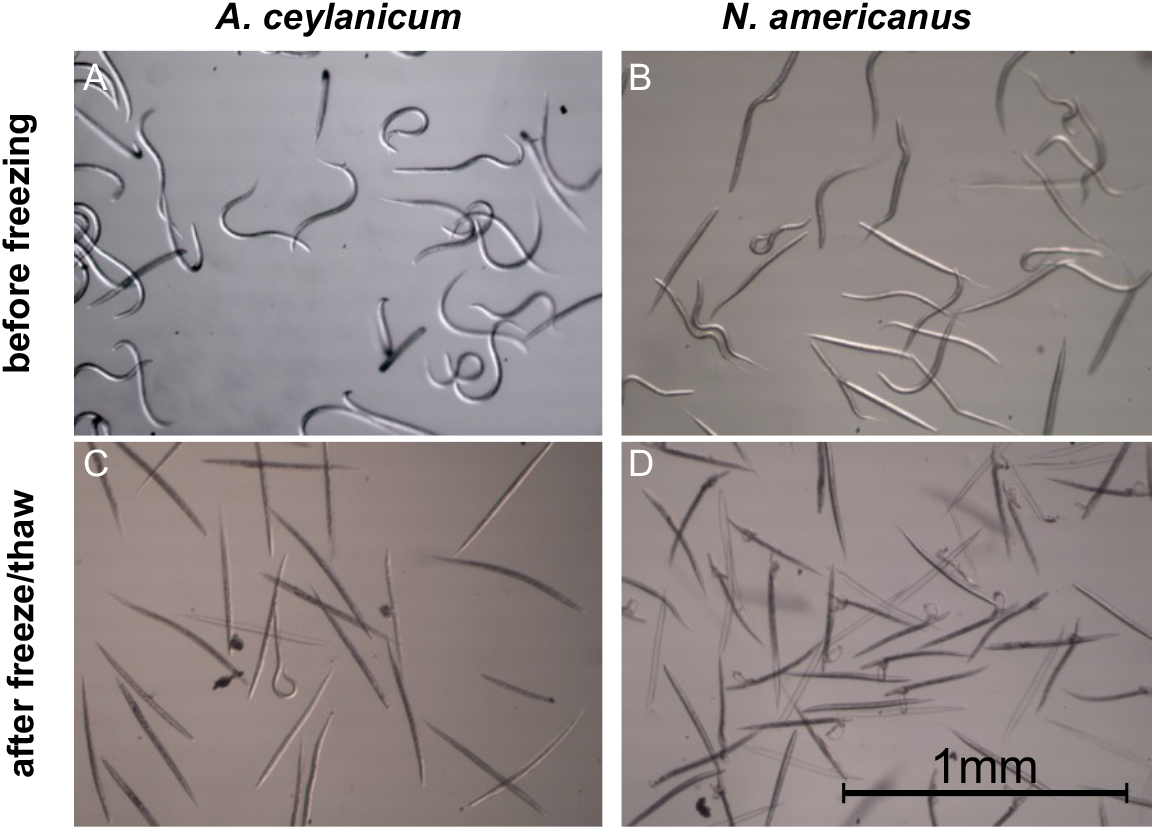
Morphology and survival of infectious third-staged larvae (iL3) from two hookworm species before and after freezing. Morphology of *A. ceylanicum* (A) and *N. americanus* (B) iL3 within one week of harvesting from charcoal culturing of infected hamster feces. The iL3 are curved, active, and alive. Morphology of thawed (C) *A. ceylanicum* and (D) *N. americanus* iL3 frozen for 2 weeks. Almost all (∼99%) of the *A. ceylanicum* iL3 and all of the *N. americanus* L3i are rigid rods— immotile and dead. Scale bar is the same in all panels.

We nonetheless tested infectivity of these cryopreserved *A. ceylanicum* iL3 hookworms. Hamsters were infected with live iL3 from either fresh charcoal cultures or recovered from cryopreserved iL3s frozen for 2 weeks or 3 months. Based both on intestinal hookworm burdens and fecal egg counts, infectivity of cryopreserved iL3s was very poor compared to infection of iL3s from fresh fecal cultures even when equal numbers of live iL3s were used for the infection (Figure 2). Infecting with iL3s frozen for 2 weeks or 3 months, *A. ceylanicum* hookworm burdens were reduced 92% and 96% respectively relative to levels found infecting with fresh iL3s; fecal egg counts were reduced 90% and 94% respectively. Because we did not recover any live *Necator* hookworms, we did not perform similar experiments with this parasite.

**Figure 2.**
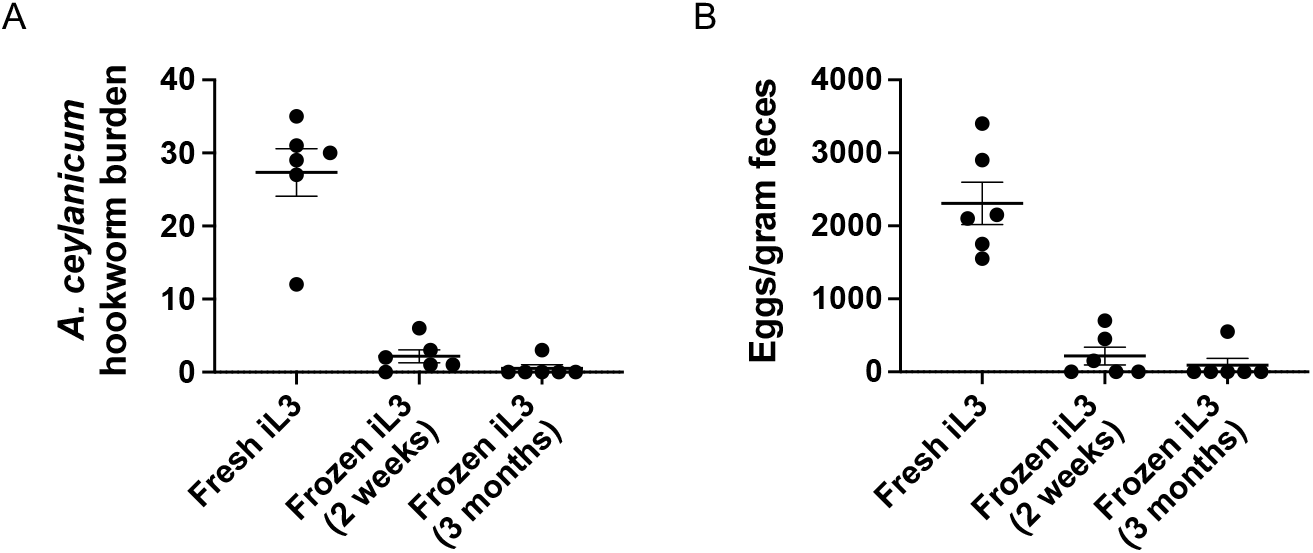
Infection efficacy of thawed *A. ceylanicum* previously frozen at iL3 stage. Average small intestinal burdens (A) and parasite fecal egg counts (B) of hamsters infected with iL3 either harvested from a fresh infection or from frozen stocks held at - 80°C for the time indicated. Each dot represents an individual hamster (n=6/group). Error bars here and elsewhere are standard error of the mean.

### Development of a method for recovering cryopreserved L1s to iL3s

Because the literature suggested cryopreservation of L1s might be superior to that of iL3s (Nolan et al. 1994), we decided to adapt the L1 method, changing the way the L1s were allowed to progress to the iL3 infectious stage. We hypothesized that growth of larvae on *E. coli* in culture might be less than ideal and that growth under conditions more similar to those used to normally give rise to iL3s in the laboratory might work better.

Hookworm eggs (*A. ceylanicum* and *N. americanus*) were purified from an overnight fecal collection of infected hamsters using standard techniques (see Materials and Methods; (Mes, Eysker, and Ploeger 2007)). After the final wash, the eggs were allowed to hatch as L1s in *Caenorhabditis elegans* S Medium (Sulston and Hodgkin 1988) at 28°C for 42 hours. The larvae (L1) were then resuspended in previously established cryoprotectant solution (Nolan et al. 1994; Duarte, Harrison, and Cappello 2003) in cryovials. The cryovials were then placed overnight in a -80°C incubator and allowed to cool at a rate of 1°C per minute using a freezing container. The following day, the cryovials were transferred to liquid nitrogen. This freezing protocol worked well based on a comparison of freshly isolated L1s from feces (Figures 3A, B, respectively *A. ceylanicum* and *N. americanus*) to freeze-thawed L1s (Figures 3C, D, respectively *A. ceylanicum* and *N. americanus*). One to three years after storage at -80° C, a high percentage of viable L1s can still be recovered for both parasites with this protocol (Figure 3; 48% and 46% respectively for *A. ceylanicum* and *N. americanus*).

**Figure 3.**
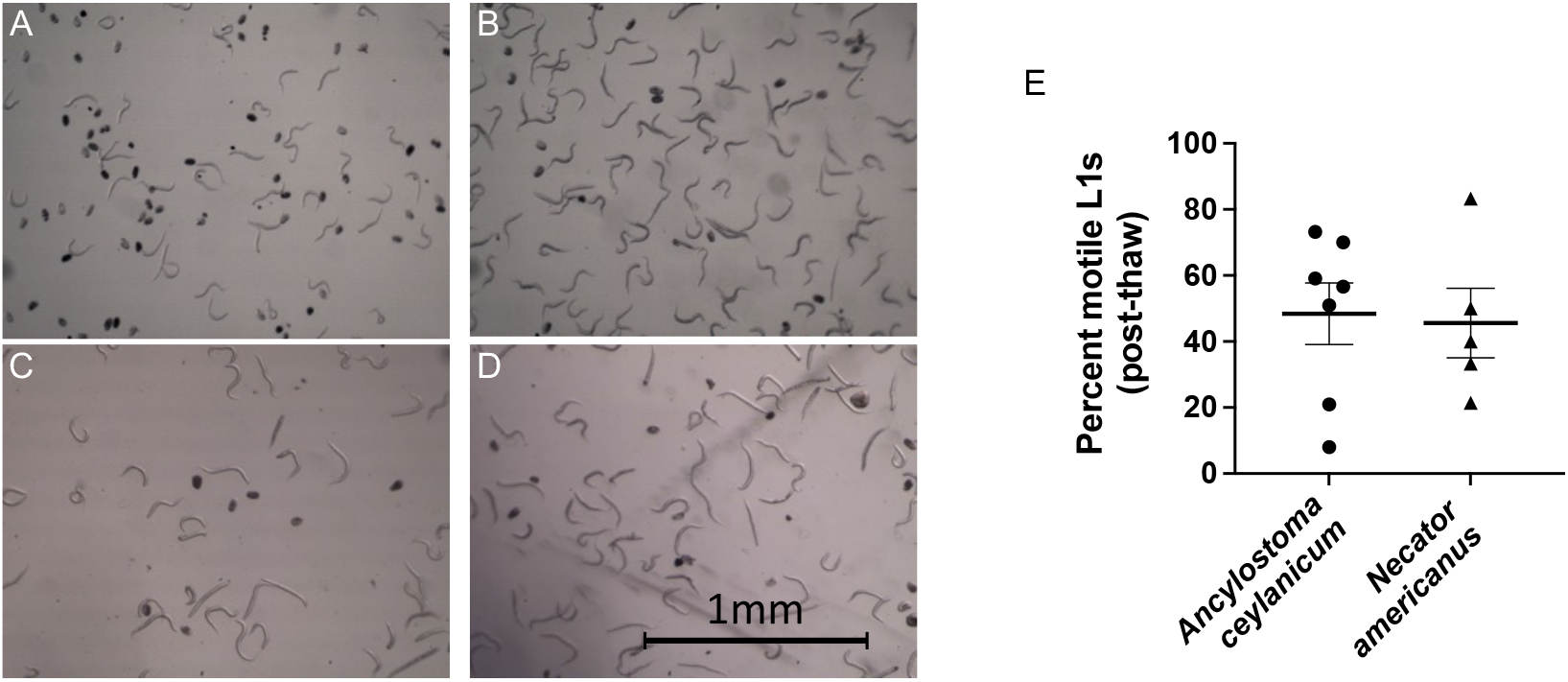
Morphology and survival of first-staged larvae (L1s) from two hookworm species before and after freezing. Morphology of *A. ceylanicum* (A) and *N. americanus* (B) L1s 42-48 hours following hatch-off of parasite eggs freshly isolated from infected feces. Many of the L1s are curved, active, and alive. Morphology of thawed (C) *A. ceylanicum* and (D) *N. americanus* L1s frozen for 15 and 36 months, respectively. Frozen L1s were thawed quickly in 55°C water bath, brought up to 50 mL volume with sterile water, spun down, resuspended in sterile water, allowed to recover for 1-2 hours at 28°C, and then imaged Many of the L1s are curved, active, and alive. Scale bar is the same in all panels. (E). The percent of thawed L1s that were alive are graphed for each hookworm species frozen for the time indicated in panels (C) and (D). An aliquot of L1’s were taken (∼100 worms per aliquot) and the motility of all the worms was noted. Each individual point is an independent thaw from independent frozen stocks (each stock was sampled three times and an average was taken).

To mimic a more natural progression to the iL3 stage than previous protocols, we thawed frozen L1s and plated them on a fecal-charcoal mixture typical for normal life cycle maintenance in the laboratory. However, unlike normal laboratory maintenance whereby feces from infected hamsters are mixed with activated charcoal and allowed to develop, here we mixed thawed L1s and added them to activated charcoal pre-mixed with *uninfected* hamster feces, providing the thawed L1s with a robust more natural fecal environment to develop in. After 7 days of larval develop on activated charcoal, the iL3s were recovered by the Baermann technique. Using this technique we found that for *A. ceylanicum and N. americanus* respectively, 31% and 16% of the total L1s (independent of whether alive or dead) plated on charcoal developed to the iL3 stage (Figure 4). Given that ∼50% of the L1s appeared alive (Figure 3), then ∼30-60% of the live L1s were able to develop to the iL3 stage using these culture conditions.

**Figure 4.**
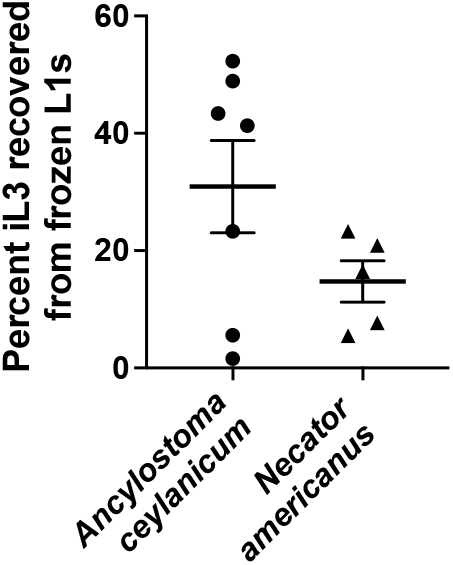
Percent iL3s recovered from frozen L1s. About 10,000 thawed L1s (without discriminating live vs dead) were pipetted onto charcoal-uninfected hamster fecal mixture, incubated at 28°C, harvested using a Baermann funnel, and then the total number of iL3s were counted. Each data point is from an independent frozen sample. The *A. ceylanicum* samples were frozen for a minimum of 1 year. The *N. americanus* samples were frozen for a minimum of two years.

### iL3s recovered from L1s frozen for up to three years are able to regenerate the hookworm life cycle

*A. ceylanicum* iL3s recovered using fecal-charcoal plates above were tested for their ability to complete the life cycle in hamsters (Figure 5). The infectivity of iL3s from an active infection (non-frozen), iL3s recovered from L1s frozen for 15 months, and iL3s recovered from L1s frozen for 36 months were used to infect hamsters. Based on intestinal hookworm burdens, there was no difference in the infectivity of iL3s from fresh L1 cultures, from 15-month frozen L1 cultures 15 months), or from 36-month frozen L1 cultures (Figure 5A; P=0.77 and 0.27, respectively, using one-way ANOVA and Dunnett’s posttest comparing each frozen group to fresh iL3s). Similarly, based on fecal egg counts, there was no difference whether infecting with iL3s from fresh L1 cultures, from 15-month frozen L1 cultures 15 months), or from 36-month frozen L1 cultures (Figure 5B; P=0.76 and 0.20, respectively, using one-way ANOVA and Dunnett’s posttest comparing each frozen group to fresh iL3s).

**Figure 5.**
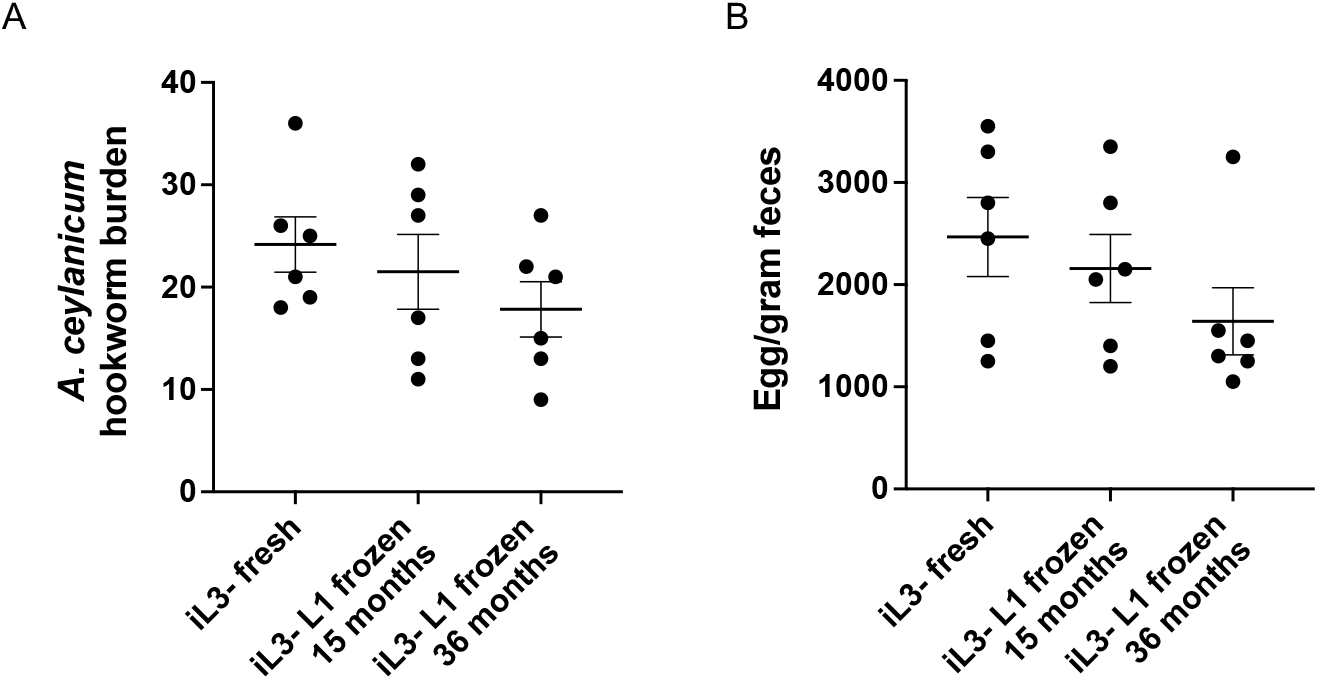
Infectivity of *A. ceylanicum* iL3s recovered from frozen L1 samples. (A). Total intestinal *A. ceylanicum* hookworm burdens of hamsters infected with iL3s recovered from charcoal plates seeded with feces from on-going infection (fresh) or from charcoal-fecal plates were seeded with cryopreseved L1s frozen for the amount of time indicated. The average worm burdens were 24.2, 21.5, and 17.8 for fresh iL3, 15-month frozen iL3s, and 36-month frozen iL3s, respectively. (B). Eggs per gram of feces from the same animals in panel A. The average eggs per gram of feces were 2467, 2158, and 1642 for fresh iL3, 15-month frozen iL3s, and 36-month frozen iL3s, respectively.

We also compared the infectivity of *N. americanus* iL3s from fresh L1s (active infection) and from L1s frozen for 36 months. In both cases, hookworm adults were found in the small intestinal tract of all hamsters, although there were noticeably fewer hookworms when infecting with iL3s from frozen L1 cultures (Figure 6A; 65% reduction; P=0.014 using student’s T-test). In both cases, parasite eggs were also found in the feces of all hamsters, although there were noticeably fewer hookworm eggs parasite eggs when infecting with iL3s from frozen L1 cultures (Figure 6B; 72% reduction; P=0.0038 using student’s T-test). Nonetheless, the infection using iL3s from L1s frozen for three years was robust.

**Figure 6.**
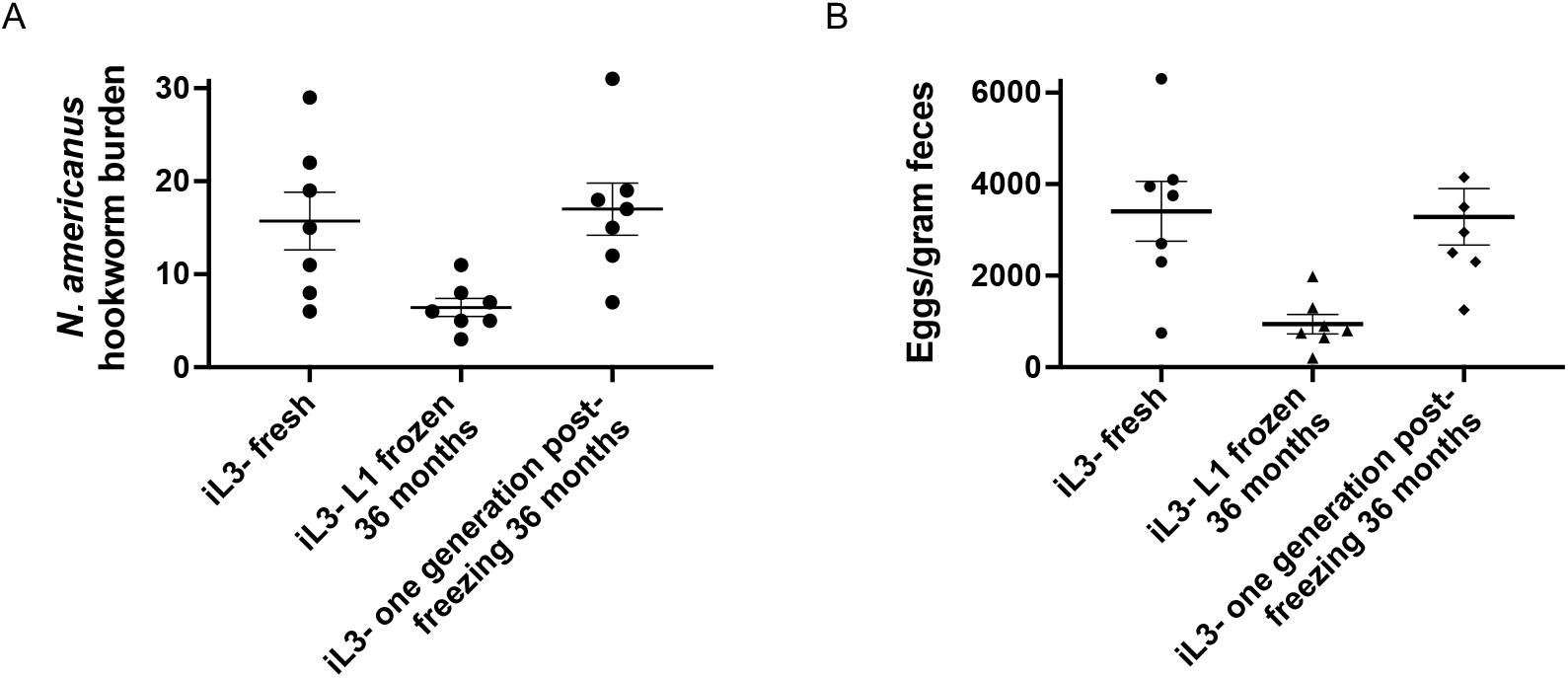
Infectivity of *N. americanus* iL3s recovered from frozen L1 samples. (A). Total intestinal *N. americanus* hookworm burdens of hamsters infected with iL3s recovered from charcoal plates seeded with feces from on-going infection (fresh), from charcoal-fecal plates were seeded with cryopreseved L1s frozen for 36 months, or from charcoal plates seeded with feces from the first generation following recovery from cryopreservation. The experiment on the right side of the graph was done at a separate time from the other two. The average worm burdens from left to right were 15.7, 6.4, and 17.0. (B). Eggs per gram of feces from the same animals in panel A. The average eggs per gram of feces were from left to right 3407, 940, and 3286.

Strikingly, we harvested parasite eggs are harvested from hamsters infected with iL3s generated from frozen L1s. When these first-generation eggs were allowed to progress to the iL3 stage and used to infect hamsters, infectivity was normal (Figure 6A, B). Thus, we were able to fully regenerate a healthy *N. americanus* lifecycle even after three years of cryopreservation.

### The protocol can be generalized to *Strongyloides ratti* and *Heligmosomoides polygyrus bakeri*

We maintain *S. ratti* in laboratory rats. The *S. ratti* lifecycle in rats is similar to that of *N. americanus* in hamsters. Feces from infected rodent hosts containing parasite eggs can be cultured on activated charcoal until the iL3 stage, at which point they can be administered subcutaneously into naive rodent hosts, allowing for completion of the life cycle. Although *S. ratti* and *N. americanus* are phylogenetically distant and appear in different clades of phylum Nematoda (clade IV and clade V, respectively; (Parkinson et al. 2004)), we tested whether or not the cryoprotection protocol could be applied to *S. ratti. S. ratti* L1s were similarly frozen as for hookworms and thawed after two weeks. Compared to L1s freshly isolated from feces (Figure 7A), L1s recovered from frozen culture appeared healthy (Figure B). Upon thawing, L1s were grown on activated charcoal mixed with uninfected rat feces until the iL3 stage. Compared to iL3s from a fresh infection cycle (Figure 7C), iL3s from derived from frozen L1s were also very healthy (Figure 7D). On average, 73±5% of thawed L1s appeared alive (motile) and 24±4% of L1s plated on fecal-charcoal mixture developed to iL3. When introduced into rats, the infectivity of iL3s cultured from frozen L1s was similar to that from iL3s cultured from fresh L1s based on fecal egg counts (Figure 7E; P=0.78).

**Figure 7.**
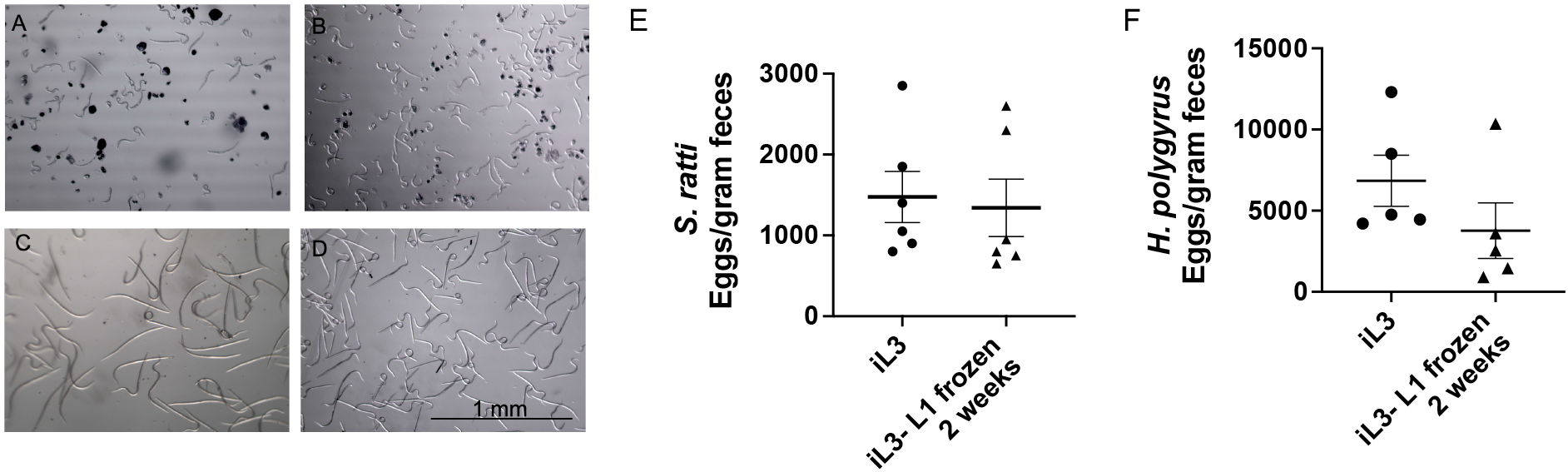
Successful cryoprotection of *S. ratti* and *H. polygyrus*. (A). Morphology of *S. ratti* L1s following hatch-off in S medium of parasite eggs freshly isolated from infected feces. (B). Morphology of *S. ratti* L1s thawed from stocks frozen for two weeks. Many of the L1s are curved, motile, and alive. (C). Morphology of *S. ratti* iL3s harvested from charcoal culture seeded with feces from an active infection. (D). Morphology of *S. ratti* iL3s cultured from L1s frozen for two weeks on charcoal seeded with rat feces. Scale bar is the same in all panels. (E). Infectivity of *S. ratti* iL3 isolated from charcoal culture plates seeded with fresh L1s (iL3) or frozen L1s based on fecal egg counts. The average fecal egg counts were 1475 and 1342 eggs per gram of feces, respectively. F. Infectivity of *H. polygyrus bakeris* iL3s isolated from charcoal culture plates seeded with fresh L1s (iL3) or frozen L1s based on fecal egg counts. The average fecal egg counts were 6840 and 3770 eggs per gram of feces, respectively. Statistically there is no difference between the two conditions (P=0.22 student’s t-test).

We also maintain *H. polygyrus bakeri* in the laboratory. This natural parasite of mice is perhaps the most common model for immunological studies of gastrointestinal nematode parasites – mammalian host interaction (Reynolds, Filbey, and Maizels 2012). Similar to *S. ratti*, we were able to successfully infect with *H. polygyrus bakeri* iL3s derived from L1s frozen for two weeks and cultured on activated charcoal mixed with uninfected mouse feces (Figure 7F).

## 4. DISCUSSION

We have demonstrated a robust method for cryopreservation of iL3s from both general of human hookworms (*A. ceylanicum, N. americanus*), the threadworm *S. ratti*, and the model gastrointestinal nematode parasite *H. polygyrus bakeri*. For hookworms, successful recovery and reintroduction of parasite into host hamsters was demonstrated for at least three years after freezing. To our knowledge, this is the first successful demonstration of freezing and recovery for this length of time and for these nematodes.

The key to this technique is the culturing of thawed (previously-frozen) L1 parasites in charcoal-fecal mixture in which the feces comes from uninfected hosts (hamsters for hookworms, rats for *S. ratti*, and mice for *H. polygyrus bakeri*). The growth of L1 to iL3 under these conditions is more robust and closer to the natural cycle than, for example, growth on a monoculture of bacteria, *e*.*g*., (Nolan et al. 1994). Conversely, our attempts at infection with of frozen iL3s resulted in poor infectivity for *A. ceylanicum* and no recovery of live hookworms for *N. americanus*.

Successful cryoprotection and recovery of gastrointestinal nematode parasites should provide a significant boost to the field of gastrointestinal nematode studies. There are significant challenges associated with studies of these parasites. Currently, most of these parasites have to be continually maintained in mammalian hosts, which is a significant burden. This burden can be additionally increased because of external events, *e*.*g*., as many laboratories discovered during COVID. Frozen stocks provide an important sense of security and obviate the need to continually maintain cultures during times where that is difficult.

Helminths parasites such as hookworms can also show great strain variability at the genome level, whereas the populations maintained in the laboratory are relatively small. Thus, we have found that sometimes laboratory populations can bottleneck and become unhealthy and crash. In these instances, frozen stocks can allow for successful recovery of such lines. Genotypes can also change over time and it may be desirable to preserve a population at a current genotype, which can be returned to in the future with frozen stocks. As hookworms for immunotherapies move towards GMP production, methods for long-term preservation of hookworms is also critically important for continued human experimental infection and therapy.

In summary, we provide here a detailed and validated methodology for long-term preservation of human hookworm lines and other gastrointestinal nematode parasites that should greatly facilitate and result in significant expansion of research in vital studies of these parasites.

## ACKNOWLEDGEMENTS

This work was financially supported by the National Institutes of Health--National Institute of Allergy and Infectious Diseases grants R01-AI056189, R01-AI150866, and R21-AI149037 and by the National Institutes of Health—Eunice Kennedy Shriver National Institute of Child Health & Human Development grant 1R01-HD099072 to R.V.A.

## REFERENCES

Aikens, L. M., and G. A. Schad. 1989. “Radiolabeling of Infective Third-Stage Larvae of Strongyloides Stercoralis by Feeding [75 Se]Selenomethionine-Labeled Escherichia Coli to First- and Second-Stage Larvae.” The Journal of Parasitology 75 (5): 735–39.

Bartsch, Sarah M., Peter J. Hotez, Lindsey Asti, Kristina M. Zapf, Maria Elena Bottazzi, David J. Diemert, and Bruce Y. Lee. 2016. “The Global Economic and Health Burden of Human Hookworm Infection.” PLoS Neglected Tropical Diseases 10 (9): e0004922.

Chapman, Paul R., Paul Giacomin, Alex Loukas, and James S. McCarthy. 2021. “Experimental Human Hookworm Infection: A Narrative Historical Review.” PLoS Neglected Tropical Diseases 15 (12): e0009908.

Chylinski, C., J. Cortet, G. Sallé, P. Jacquiet, and J. Cabaret. 2015. “Storage of Gastrointestinal Nematode Infective Larvae for Species Preservation and Experimental Infections.” Parasitology Research 114 (2): 715–20.

Duarte, John, Lisa M. Harrison, and Michael Cappello. 2003. “Short Report: Ancylostoma Ceylanicum: Exsheathment Is Not Required for Successful Cryopreservation of Third Stage Hookworm Larvae.” The American Journal of Tropical Medicine and Hygiene 68 (1): 44–45.

Fumagalli, Matteo, Uberto Pozzoli, Rachele Cagliani, Giacomo P. Comi, Nereo Bresolin, Mario Clerici, and Manuela Sironi. 2010. “The Landscape of Human Genes Involved in the Immune Response to Parasitic Worms.” BMC Evolutionary Biology 10 (1): 264.

Fumagalli, Matteo, Manuela Sironi, Uberto Pozzoli, Anna Ferrer-Admetlla, Linda Pattini, and Rasmus Nielsen. 2011. “Signatures of Environmental Genetic Adaptation Pinpoint Pathogens as the Main Selective Pressure through Human Evolution.” PLoS Genetics 7 (11): e1002355.

Haldeman, Matthew S., Melissa S. Nolan, and Kija R. N. Ng’habi. 2020. “Human Hookworm Infection: Is Effective Control Possible? A Review of Hookworm Control Efforts and Future Directions.” Acta Tropica 201 (January): 105214.

Hamory, Joan, Edward Miguel, Michael Walker, Michael Kremer, and Sarah Baird. 2021. “Twenty-Year Economic Impacts of Deworming.” Proceedings of the National Academy of Sciences of the United States of America 118 (14): e2023185118.

Hu, Yan, Sophia B. Georghiou, Alan J. Kelleher, and Raffi V. Aroian. 2010. “Bacillus Thuringiensis Cry5B Protein Is Highly Efficacious as a Single-Dose Therapy against an Intestinal Roundworm Infection in Mice.” PLoS Neglected Tropical Diseases 4 (3): e614.

Hu, Yan, Melanie M. Miller, Alan I. Derman, Brian L. Ellis, Rose Gomes Monnerat, Joe Pogliano, and Raffi V. Aroian. 2013. “Bacillus Subtilis Strain Engineered for Treatment of Soil-Transmitted Helminth Diseases.” Applied and Environmental Microbiology 79 (18): 5527–32.

Hu, Yan, Melanie Miller, Bo Zhang, Thanh-Thanh Nguyen, Martin K. Nielsen, and Raffi V. Aroian. 2018. “In Vivo and in Vitro Studies of Cry5B and Nicotinic Acetylcholine Receptor Agonist Anthelmintics Reveal a Powerful and Unique Combination Therapy against Intestinal Nematode Parasites.” PLoS Neglected Tropical Diseases 12 (5): e0006506.

Hu, Yan, Thanh-Thanh Nguyen, Alice C. Y. Lee, Joseph F. Urban Jr, Melanie M. Miller, Bin Zhan, David J. Koch, et al. 2018. “Bacillus Thuringiensis Cry5B Protein as a New Pan-Hookworm Cure.” International Journal for Parasitology, Drugs and Drug Resistance 8 (2): 287–94.

Hu, Yan, Bin Zhan, Brian Keegan, Ying Y. Yiu, Melanie M. Miller, Kathryn Jones, and Raffi V. Aroian. 2012. “Mechanistic and Single-Dose in Vivo Therapeutic Studies of Cry5B Anthelmintic Action against Hookworms.” PLoS Neglected Tropical Diseases 6 (11): e1900.

Jimenez Castro, Pablo D., Abdelmoneim Mansour, Samuel Charles, Joe Hostetler, Terry Settje, Daniel Kulke, and Ray M. Kaplan. 2020. “Efficacy Evaluation of Anthelmintic Products against an Infection with the Canine Hookworm (Ancylostoma Caninum) Isolate Worthy 4.1F3P in Dogs.” International Journal for Parasitology, Drugs and Drug Resistance 13 (August): 22–27.

Jimenez Castro, Pablo D., Abhinaya Venkatesan, Elizabeth Redman, Rebecca Chen, Abigail Malatesta, Hannah Huff, Daniel A. Zuluaga Salazar, Russell Avramenko, John S. Gilleard, and Ray M. Kaplan. 2021. “Multiple Drug Resistance in Hookworms Infecting Greyhound Dogs in the USA.” International Journal for Parasitology, Drugs and Drug Resistance 17 (December): 107–17.

Jourdan, Peter Mark, Poppy H. L. Lamberton, Alan Fenwick, and David G. Addiss. 2018. “Soil-Transmitted Helminth Infections.” The Lancet 391 (10117): 252–65.

Li, Hanchen, Ambily Abraham, David Gazzola, Yan Hu, Gillian Beamer, Kelly Flanagan, Ernesto Soto, et al. 2021. “Recombinant Paraprobiotics as a New Paradigm for Treating Gastrointestinal Nematode Parasites of Humans.” Antimicrobial Agents and Chemotherapy 65 (3). https://doi.org/10.1128/AAC.01469-20.

Loukas, Alex, Peter J. Hotez, David Diemert, Maria Yazdanbakhsh, James S. McCarthy, Rodrigo Correa-Oliveira, John Croese, and Jeffrey M. Bethony. 2016. “Hookworm Infection.” Nature Reviews. Disease Primers 2 (December): 16088.

Loukas, Alex, Rick M. Maizels, and Peter J. Hotez. 2021. “The Yin and Yang of Human Soil-Transmitted Helminth Infections.” International Journal for Parasitology 51 (13–14): 1243–53.

Maizels, Rick M., Edward J. Pearce, David Artis, Maria Yazdanbakhsh, and Thomas A. Wynn. 2009. “Regulation of Pathogenesis and Immunity in Helminth Infections.” The Journal of Experimental Medicine 206 (10): 2059–66.

Mes, Ted H. M., Maarten Eysker, and Harm W. Ploeger. 2007. “A Simple, Robust and Semi-Automated Parasite Egg Isolation Protocol.” Nature Protocols 2 (3): 486–89.

Montaño, Karen J., Carmen Cuéllar, and Javier Sotillo. 2021. “Rodent Models for the Study of Soil-Transmitted Helminths: A Proteomics Approach.” Frontiers in Cellular and Infection Microbiology 11 (April): 639573.

Mourão Dias Magalhães, Luisa, Livia Silva Araújo Passos, Ricardo Toshio Fujiwara, and Lilian Lacerda Bueno. 2021. “Immunopathology and Modulation Induced by Hookworms: From Understanding to Intervention.” Parasite Immunology 43 (2): e12798.

Nolan, T. J., V. M. Bhopale, Z. Megyeri, and G. A. Schad. 1994. “Cryopreservation of Human Hookworms.” The Journal of Parasitology 80 (4): 648–50.

Noon, Jason B., Erich M. Schwarz, Gary R. Ostroff, and Raffi V. Aroian. 2019. “A Highly Expressed Intestinal Cysteine Protease of Ancylostoma Ceylanicum Protects Vaccinated Hamsters from Hookworm Infection.” PLOS Neglected Tropical Diseases. https://doi.org/10.1371/journal.pntd.0007345.

Parkinson, John, Makedonka Mitreva, Claire Whitton, Marian Thomson, Jennifer Daub, John Martin, Ralf Schmid, et al. 2004. “A Transcriptomic Analysis of the Phylum Nematoda.” Nature Genetics 36 (12): 1259–67.

Pullan, Rachel L., Jennifer L. Smith, Rashmi Jasrasaria, and Simon J. Brooker. 2014. “Global Numbers of Infection and Disease Burden of Soil Transmitted Helminth Infections in 2010.” Parasites & Vectors 7 (January): 37.

Reynolds, Lisa A., Kara J. Filbey, and Rick M. Maizels. 2012. “Immunity to the Model Intestinal Helminth Parasite Heligmosomoides Polygyrus.” Seminars in Immunopathology 34 (6): 829–46.

Ryan, Stephanie M., Ramon M. Eichenberger, Roland Ruscher, Paul R. Giacomin, and Alex Loukas. 2020. “Harnessing Helminth-Driven Immunoregulation in the Search for Novel Therapeutic Modalities.” PLoS Pathogens 16 (5): e1008508.

Shepherd, Catherine, Phurpa Wangchuk, and Alex Loukas. 2018. “Of Dogs and Hookworms: Man’s Best Friend and His Parasites as a Model for Translational Biomedical Research.” Parasites & Vectors 11 (1): 59.

Sulston, J., and J. Hodgkin. 1988. “Methods.” In The Nematode Caenorhabditis Elegans, edited by W. B. Wood, 587–606. New York, NY: Cold Spring Harbor Laboratory Press.

Umbrello, G., R. Pinzani, A. Bandera, F. Formenti, G. Zavarise, M. Arghittu, D. Girelli, et al. 2021. “Hookworm Infection in Infants: A Case Report and Review of Literature.” Italian Journal of Pediatrics 47 (1): 26.

